# Prediction of kinase-specific phosphorylation sites through an integrative model of protein context and sequence

**DOI:** 10.1101/043679

**Authors:** Ralph Patrick, Coralie Horin, Bostjan Kobe, Kim-Anh Lê Cao, Mikael Bodén

**Affiliations:** School of Chemistry and Molecular Biosciences, The University of Queensland, St Lucia, 4072, Australia; Polytech Nice-Sophia, Université Nice Sophia-Antipolis, Nice, 06103, France; Institute for Molecular Bioscience, The University of Queensland, St Lucia, 4072, Australia; Australian Infectious Diseases Research Centre, The University of Queensland, St Lucia, 4072, Australia; The University of Queensland Diamantina Institute, Translational Research Institute, Woolloongabba, QLD, 4102, Australia

## Abstract

The identification of kinase substrates and the specific phosphorylation sites they regulate is an important factor in understanding protein function regulation and signalling pathways. Computational prediction of kinase targets – assigning kinases to putative substrates, and selecting from protein sequence the sites that kinases can phosphorylate – requires the consideration of both the cellular context that kinases operate in, as well as their binding affinity. This consideration enables investigation of how phosphorylation influences a range of biological processes.

We report here a novel probabilistic model for the classification of kinase-specific phosphorylation sites from sequence across three model organisms: human, mouse and yeast. The model incorporates position-specific amino acid frequencies, and counts of co-occurring amino acids from kinase binding sites in a kinase‐ and family-specific manner. We show how this model can be seamlessly integrated with protein interactions and cell-cycle abundance profiles. When evaluating the prediction accuracy of our method, PhosphoPICK, on an independent hold-out set of kinase-specific phosphorylation sites, we found it achieved an average specificity of 97% while correctly predicting 32% of true positives. We also compared PhosphoPICK’s ability, through cross-validation, to predict kinase-specific phosphorylation sites with alternative methods, and found that at high levels of specificity PhosphoPICK outperforms alternative methods for most comparisons made.

We investigated the relationship between experimentally confirmed phosphorylation sites and predicted nuclear localisation signals by predicting the most likely kinases to be regulating the phosphorylated residues immediately upstream or downstream from the localisation signal. We show that kinases PKA, Akt1 and AurB have an over-representation of predicted binding sites at particular positions downstream from predicted nuclear localisation signals, demonstrating an important role for these kinases in regulating the nuclear import of proteins.

PhosphoPICK is freely available online as a web-service at http://bioinf.scmb.uq.edu.au/phosphopick.

## Introduction

Kinases regulate a wide variety of essential biological processes through protein phosphorylation, including transcription factor activity,^1^ the control of DNA damage repair pathways,^2^ the progression of cells through mitosis,^3^ and protein import into the nucleus.^4^ Knowledge of the kinases that regulate phosphorylation substrates is therefore a significant factor in understanding the functional consequences of protein phosphorylation events. While hundreds of thousands of phosphorylation sites have been identified across thousands of proteins,^5^ the kinases that regulate these sites in most cases remain unknown. Computational methods that predict kinase-specific phosphorylation sites are therefore an important contributor to understanding the role of phosphorylation events in biological processes.^6^ Such methods contribute to the guidance of phosphorylation experiments^7^ and provide information about the likely signalling pathways that phosphorylation sites may be involved in.^8^

Kinase-mediated phosphorylation is regulated by several important factors that can be leveraged to build predictive models. One is the sequence-level motifs surrounding phosphorylation sites that interact with kinase binding domains. The protein sequence determines whether a kinase can bind to the protein; previous studies have shown that local motifs surrounding a phosphorylation site interact with the binding domain of kinases to allow phosphorylation.^9,10^ There are numerous kinase-specific phosphorylation site predictors that take advantage of the sequence specificity of kinases to predict kinase-specific phosphorylation sites^11–13^ as well as phosphorylation sites in a non-kinase specific manner.^14,15^

The presence of valid kinase-binding motifs on a protein is no guarantee that a kinase will phosphorylate a substrate however.^16^ The targeting of phosphorylation substrates by kinases is subject to, and controlled by, a wide variety of processes within the cell – what may be called the “context factors” that ensure kinase-substrate fidelity. Context factors can include proteins that mediate the interaction between kinases and their substrates,^17^ activating proteins such as cyclins,^18^ sub-cellular compartmentalisation^19^ and the various stages within the mitotic cell cycle.^20^

We have shown previously that context information (in the form of protein-protein interaction and association data, as well as protein abundance levels across the cell cycle) can be incorporated into a probabilistic model that maps kinases to putative substrates.^21^ This model not only provides an accurate predictor of kinase substrates, but importantly, the sequence-level prediction of kinase-specific phosphorylation sites can be greatly enhanced by the model’s additional predictive power. While this model was able to use context alone to predict kinase substrates, we hypothesised that the incorporation of sequence and context into a single model would provide better explanatory power of the factors that describe kinase targets.

In this paper, we present a novel probabilistic method for predicting kinase-specific phosphorylation sites that incorporates position-specific amino acid frequencies and counts of co-occurring neighbouring amino acids in a family-specific manner across three model organisms: human, mouse and yeast. We demonstrate that this sequence model can be used as a module within a larger Bayesian network that describes the context factors that influence how a kinase targets a protein substrate. The seamless integration of these two domains of information – context and sequence – allows for a comprehensive model of kinase-protein phosphorylation. We compare the ability of our method, PhosphoPICK, to predict kinase-specific phosphorylation sites against alternative phosphorylation predictors, and show that PhosphoPICK has a superior ability to predict kinase-specific phosphorylation sites for most comparisons made.

As we now have a predictor that ably integrates the context and sequence conditions that regulate phosphorylation, we are in a position to investigate phosphorylation-dependent functions and probe the kinases that are involved in regulating these functions. The nuclear import of proteins is a highly-specific process, involving the binding of importin proteins to cargo proteins that contain a nuclear localisation signal (NLS).^22,23^ It has been shown that the binding of importin proteins to their cargo can be promoted or inhibited by the presence of phosphorylation adjacent to the NLS.^24^ We therefore investigated the relationship between nuclear localisation signals and phosphorylation by cross-referencing experimentally identified phosphorylation sites with predicted NLSs. We used PhosphoPICK to identify the most likely candidate kinases for NLS-adjacent phosphorylation sites, and performed a statistical analysis to identify sites relative to NLSs that have an over-representation of kinase binding sites. We identify several kinases as candidates to regulate phosphorylation sites at sites downstream from the NLSs, most notably protein kinase A (PKA), Akt1 and Aurora kinase B (AurB). We also identify kinases that regulate sites upstream from the NLS, including cyclin dependent kinase 2 (CDK2). Gene ontology (GO) term enrichment analyses indicate that the phosphorylation of specific sites close to the NLS by these kinases regulate distinct biological functions.

## Experimental procedures

### Data resources

We obtained kinase-specific phosphorylation data for human and mouse from PhosphoSitePlus^®^, www.phosphosite.org^5^ and for yeast (Saccaromyces cerevisiae) from PhosphoGRID,^25^ which is a database of *in vivo* phosphorylation sites. For data collected from PhosphoSitePlus^®^, we ensured that phosphorylation sites used were known to occur *in vivo*, but for both databases, the kinase annotations are often informed by *in vitro* or *in vivo* experiments. We chose phosphorylation site data for kinases where there were greater than 5 unique kinase substrates, resulting in 5,209 kinase-specific phosphorylation sites across 1,826 proteins for human, 956 kinases-specific phosphorylation sites across 417 proteins for mouse, and 2,219 kinase-specific phosphorylation sites across 722 substrates for yeast. In order to have a more extensive background of phosphorylation events for training a sequence model, we also used phosphorylation sites that did not have a kinase assigned to them. We used phosphorylation sites from PhosphoSitePlus^®^ that were generated using low-throughput methods; similarly for PhosphoGRID, sites were included if they were identified using more than one method, or if the single detection method was not mass spectrometry. This resulted in an additional 5,939 phosphorylation sites for human, 2,865 additional phosphorylation sites for mouse and 674 additional phosphorylation sites for yeast.

Protein-protein interaction (PPI) data were sourced from BioGRID,^26^ protein-protein association data from STRING,^27^ and protein abundance data across the cell cycle from the work by Olsen and colleagues.^28^ As the cell-cycle information was only available for human, cell-cycle data were not incorporated into the mouse or yeast kinase models. A detailed description of how this data were curated and processed is available in. ^21^

In order to evaluate the prediction accuracy of our method on completely novel data, we created a hold-out set for kinases for which there were more than 100 known substrates – there were nine such human kinases. For each of the nine kinases, we selected a random set of substrates equal to 10% of that kinase’s substrates that were *not* in the original set of substrates used for developing the model.^21^ These substrates were excluded from all analyses and simulations, and were used only for a final evaluation of model accuracy. This resulted in a hold-out set of 145 proteins – containing 416 phosphorylation sites specific to the nine kinases. After removing the hold-out set, a set of 1,671 human proteins and 4,907 kinase-specific human phosphorylation sites remained for training and testing.

In addition, we built similarity-reduced sets of the phospho-peptide sequences obtained from PhosphoSitePlus and PhosphoGRID in order to determine whether sequence similarity could be inflating prediction accuracy. The BLASTP program^29^ was used to perform a pairwise sequence similarity comparison of each of the phospho-peptides, using 15-residue sequences centred on the phosphorylation site. All 15-residue pairs obtaining a BLASTP Evalue under 0.05, with sequence identity of at least 30%, were retained. Similar pairs within the same kinase category were reduced through the arbitrary removal of one of the phospho-peptides; phospho-peptides that were similar, but phosphorylated by different kinases, were not reduced. The similarity reduction was also applied to the background set of peptides.

### PhosphoPICK method and workflow

Building on our existing context model, we developed a model for predicting kinase-specific phosphorylation sites from sequence, as well as a model that incorporates this sequence model into the context model described in our previous work. The data used for training the models are available in Data S1.

#### Sequence model

We present a Bayesian network model for modelling various sequence features of a kinase binding motif (Fig. 1(a)). We represent potential amino acid residues in an *n* length sequence motif surrounding a phosphorylation site as discrete variables conditioned on two Boolean variables. The first represents the event that some kinase of interest, *K*, binds to the site, the second represents the event that a family member (i.e. any family member of *K*) binds to the site. Each variable – *R*_–*m*_ to *R*_+*m*_, where *R*_0_ represents the site for which phosphorylation is predicted – contains three distributions of amino acid frequencies. These represent (1) the probability of each amino acid occurring at the position where *K* is seen to be phosphorylating, (2) the amino acid frequencies for binding sites from the family members of *K*, and (3) the amino acid frequency background as seen across all other phosphorylation sites in the training set.

**Figure 1:**
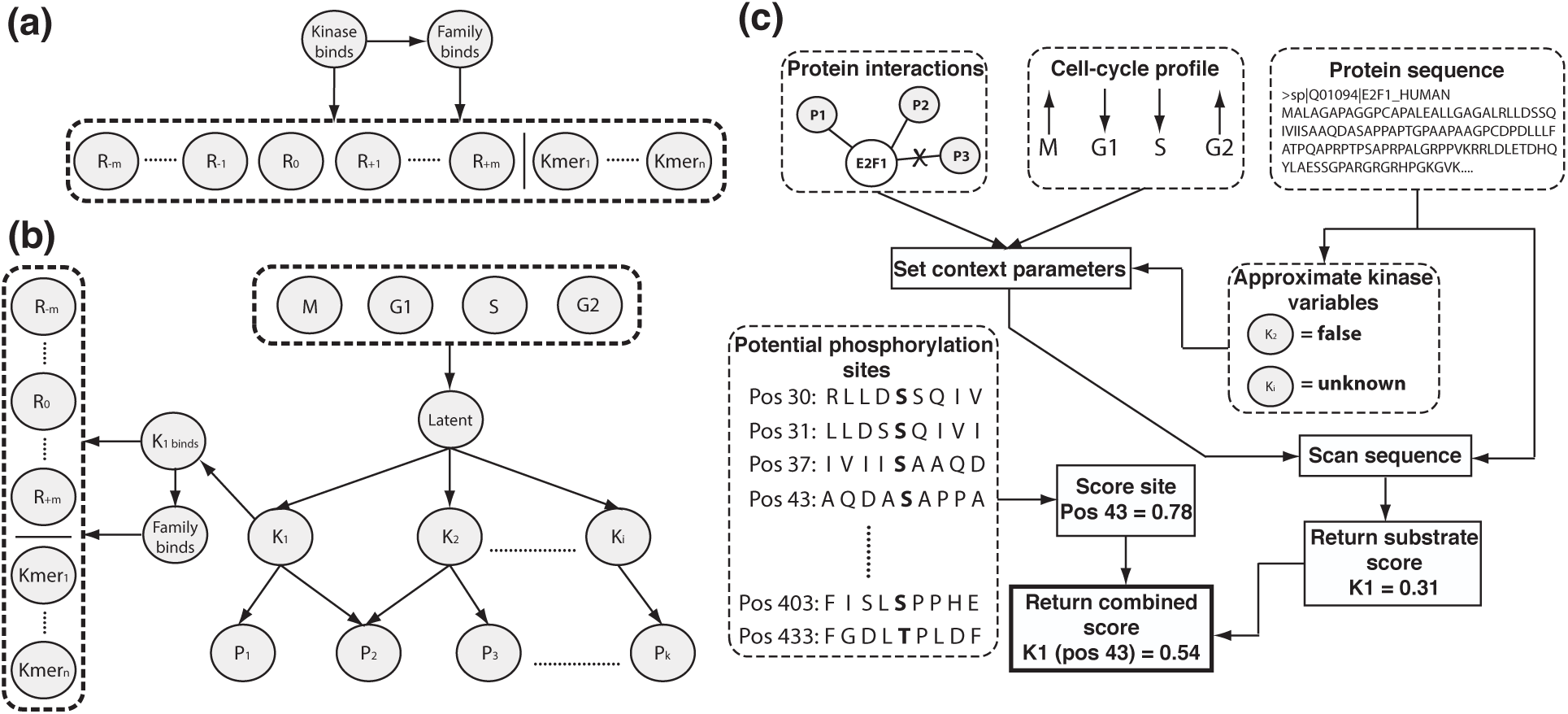
PhosphoPICK Bayesian networks and workflow. **(a)** Sequence model. *R* nodes represent positions in a motif surrounding the phosphorylation site, where *R*_0_ is the potential phosphorylation site. *Kmer*_*1*_ to *Kmer*_*n*_ represent the dimer and trimer configurations incorporated into the model. **(b)** PhosphoPICK Bayesian network model incorporating both context and sequence data. The bottom layer of nodes (*P*_1_ to *P*_k_) represent protein interactions incorporated into the model. These are conditioned on relevant kinases (*K*_1_ to *K*_*i*_), which are themselves conditioned on a latent node incorporating variables representing the four cell cycle stages. The *K*_1_*binds* “sequence” variable is conditioned on its corresponding *K*_1_ “context” variable. (c) Diagram showing the workflow involved when a kinase is queried for a protein submitted to the model. BioGRID and STRING are queried to identify what proteins the substrate interacts with, and the protein-interaction variables are set accordingly. If cell-cycle data is available, it is included also. The substrate sequence is used to estimate what kinases in the model will *not* bind to the substrate, with the remainder left unspecified. The model is then scanned across the sequence to identify the highest probability of the kinase phosphorylating the substrate. Separately, the sequence model is used to score all potential sites in the query substrate. The final prediction for a potential phosphorylation site is the average of the substrate and site score.

In addition to position-specific amino acid frequencies, we included k-mers of k=2 (dimers) and k=3 (trimers) to encode the frequency of co-occurring neighbouring amino acids. This should allow the model to capture some paired dependencies that may exist between amino acids. In order to avoid over-parameterising the sequence model with all possible combinations of dimers and trimers, we only added the k-mers that were observed in some *θ* percentage of kinase binding motifs from a training set. During cross-validation, the training set of kinase-binding motifs was taken, and k-mers observed within the motifs were counted. If a k-mer occurred in more than the *θ* percentage threshold of substrates, the k-mer was added to the model. We tested three cut-offs of *θ*: 5, 10 and 20, and found that 5 gave the best prediction accuracy across the full set of kinases (Table S-1, Table S-2 & Table S-3). As shown in Fig. 1(a), the k-mers are represented as a series of *n* Boolean variables, *Kmer*_1_ to *Kmer*_*n*_, where a k-mer is considered to be `true` if it is observed in the amino acid motif surrounding the phosphorylation site. The k-mer nodes were trained to capture the probability of each k-mer occurring within a kinase’s binding motif, that of its family members and the background set of phosphorylation sites.

It has been shown previously that varying the motif length in predicting kinase binding sites improves prediction accuracy.^13^ Therefore, for each kinase we tested five different window sizes centred around the phosphorylated residue: 7, 9, 11, 13 and 15. For each kinase we selected the window size that gave the best prediction accuracy as measured within a cross-validation test (Table S-4, Table S-5 & Table S-6).

#### Combined model

The combined model retains the structure of the “context” Bayesian network described previously,^21^ but with the sequence model incorporated into it. This model represents observations about kinase-substrate phosphorylation events, protein-protein interaction/association events believed to be relevant to kinases encoded in the model, and cell-cycle profiles of substrates as Boolean variables. A connection between a kinase and a PPI event is defined if the protein is interacting with at least 5 of the kinase’s substrates. Up to 25 connections between a kinase and a PPI event can be defined.

The sequence model was incorporated into the larger context model in a kinase-specific manner, such that for each kinase the kinase target variable in the sequence model is conditioned on the variable in the context model representing the kinase phosphorylating a substrate (Fig. 1(b)). We created models based on sets of kinases as they are classified into family similarity.^30^ For human, we created eight family-specific models comprising kinases from the CMGC (cyclin-dependent, mitogen-activated, glycogen synthase and Cdc2-like), AGC (protein kinase A, G and C families), CAMK (Ca^2+^/calmodulin-dependent kinase), TK (tyrosine kinase), “other”, STE, CK1 (cell kinase 1) and atypical kinase families. For mouse, we created three models with kinases from the CMGC, AGC and TK families; and for yeast we created four models from the CMGC, AGC, CAMK and other kinase families.

#### Setting non-query kinase nodes

The model relies partly on the expected activity of alternative kinases that are encoded in the Bayesian network. However, there is no experimental information on kinase binding events for the majority of proteins, and negative evidence (a protein *not* being phosphorylated by a particular kinase) is non-existent. Therefore we employ the amino acid sequence of a query protein to estimate what kinases in the model will not bind to the protein, and can therefore be set to `false`. In order to decide when kinase variables in the model should be set to `false`, the following steps were followed for each non-query kinase. Within a training fold, the positive training samples for that kinase were set aside. 75% of the substrates within the negative set were selected randomly, and each phosphorylation site within this set was added to the training data, while the remaining substrates were set aside as a test set.

The sequence model was then trained using the selected training samples, and used to scan over each of the substrates within the test set. The highest score for each of the substrates was recorded. The median value of these scores was then taken as a threshold representing the highest expected score for a protein that is not phosphorylated by the kinase. When evaluating the model on a test substrate, for each non-query kinase node its sequence model is used to scan the substrate and the highest score is recorded. If the score falls below the calculated threshold value, that kinase node is set to `false`, otherwise it remains unspecified.

#### Prediction workflow

A diagram illustrating the PhosphoPICK workflow for generating a prediction is shown in Fig. 1(c). To determine the probability of a query kinase phosphorylating a given substrate, the relevant context data are queried and the corresponding nodes in the Bayesian network are instantiated. Non-query kinase nodes are either set to `false` or left unspecified based on the predicted probability that the kinase can bind the substrate sequence.

The model is then scanned over the substrate’s amino acid sequence, and for every potential phosphorylation site, the n length motif corresponding to the query kinase surrounding the phosphorylation site is used to set the sequence nodes in the network. For every potential phosphorylation site, the node representing the kinase phosphorylating a substrate is queried, and the highest probability for the scan is taken as the score for that substrate.Separately, the potential phosphorylation sites within the substrate are scored using the sequence model. The final score for a kinase-specific phosphorylation site prediction is equal to the average of the substrate score from the combined model, and the site score from the sequence model.

## Model training

### Sequence model

The nodes in the sequence Bayesian network are defined using conditional probability tables (CPTs), which learn from training data all possible values that a variable can take given the set of parents it is conditioned on. If a variable does not have parents, the CPT will represent the observed frequency from the training data of it being true. As there may be amino acids or k-mers that do not occur in some of the training data, we added a uniform pseudo-count of 0.05 to all the amino acid and k-mer nodes, ensuring that the model does not consider some amino acids or k-mers impossible to occur.

### Combined model

The nodes in the combined model are defined using CPTs and our variation on the NoisyOR node,^21^ which allows for an approximation of a CPT. The protein interaction nodes were defined using NoisyOR variables, allowing parameters to be inferred even in the case of data sparsity. All other variables in the combined model were defined as CPTs.

As the combined model incorporates data representing different problems – that of predicting kinase substrates, and predicting kinase binding sites, the model was trained in two stages. First, the set of unique substrates was presented for expectation maximisation training ^31^ in order to set the parameters for the protein-interaction, cell-cycle and kinase nodes in the network. The parameters for these variables were then locked in place. Next, the sequence module within the network was trained using the set of phosphorylation sites contained in the training fold, with the position-specific amino acid nodes and k-mer nodes being set as for the sequence model. There will be some cases in the phosphorylation site data where a kinase will be phosphorylating a substrate, but not the site. In these cases, the node representing the kinase binding the substrate was set to `false`.

## Evaluating model prediction accuracy

The prediction accuracy of the models was evaluated across the 107 human kinases, 24 mouse kinases and 26 yeast kinases using ten-fold cross-validation across ten randomised data-set splits. The prediction accuracy of the sequence model was evaluated by its ability to correctly classify kinase-specific phosphorylation sites out of the set of known kinase-binding sites, and the combined model was evaluated by its ability to correctly classify kinase substrates out of the set of substrates.

To ascertain the effect that our sequence model features have on prediction accuracy, we evaluated the accuracy of a simple baseline sequence model that only contained the position-specific amino acid nodes conditioned on the kinase variable (the family variable was excluded). We also evaluated the prediction accuracy of the context model (the combined model excluding the sequence information) and compared its accuracy with the combined model to ascertain what improvement may be gained from incorporating sequence and context information into a single model. Prediction accuracy was determined using receiver operating characteristic (ROC) and calculation of area under the ROC curve (AUC) as a measure of overall model performance.^32^ We also calculated area under the ROC curve up to the fiftieth false positive (AUC50) as a measure of performance at low false-positive levels.

### Comparisons to alternative methods

We compared the ability of the complete PhosphoPICK work-flow to predict kinase-specific phosphorylation sites out of all potential phosphorylation sites in the substrate sequences. The comparison was performed firstly against the sequence model only, and secondly against three alternative methods that have a larger number of kinases available for making predictions: GPS 2.1,^13^ NetPhorest 2.0^33^ and NetworKIN 3.0.^33^ We downloaded the standalone prediction software for each of the three methods and ran the set of 1,671 proteins through them. For NetworKIN and NetPhorest, we did not specify the sites we wanted predictions for. We used GPS’s batch prediction system to run GPS on the protein set, selecting the “no threshold” option.

In order to compare PhosphoPICK predictions to the alternative methods, we again did a 10x ten-fold cross-validation run of the combined model as well as of the sequence model. As most of the potential phosphorylation sites in the substrates were not in the set of peptides used for training the sequence model (and therefore not part of the cross-validation run), the fully trained sequence model was used to score potential phosphorylation sites outside of the training set.

Due to the large number of potential phosphorylation sites being scored (~170,000 S/T sites and ~30,000 Y sites), we calculated sensitivity for two stringent levels of specificity – 99.9% and 99%. The difference in sensitivity between PhosphoPICK and each alternative was calculated across all ten cross-validation runs.

## Calculating significance of predictions

Users of the PhosphoPICK web-server are provided with an option to include empirical Pvalue calculations alongside their predictions, allowing for a measure of the significance of the predictions. To obtain empirical P-values, we first calculated proteome-wide distributions of predictions; i.e. for all kinases, substrate predictions were obtained for every protein in the relevant proteome (human, mouse or yeast), and site predictions were made for all potential phosphorylation sites in the proteome. To calculate a combined P-value for a prediction, Fisher’s method for combining probabilities was applied such that:

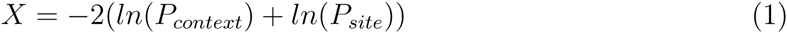

where *P*_*context*_ and *P*_*site*_ represent the P-value value calculated for a context score given to a substrate and a motif score given to a site respectively, and X follows a Chi squared distribution with 4 degrees of freedom.

### Evaluation using the hold-out set

When evaluating the performance of the model on the hold-out set, the full sets of training data were used to train the model. We predicted each potential phosphorylation site (all S/T residues for serine/threonine kinases and all Y residues for the tyrosine kinase Src) in the hold-out sequences, and evaluated the performance of the model for each kinase by its ability to predict the kinases’ phosphorylation sites out of all potential sites. In order to evaluate how well the method would be expected to perform using the P-value based thresholding system on the web-server, P-values were calculated for the predictions, and if a P-value for a prediction fell below 0.005 the prediction was considered to be `true`, and `false` otherwise.

We calculated sensitivity, specificity, balanced accuracy (BAC) and Matthews’ correlation coefficient (MCC). The metrics are defined as follows, where *TP* is the number of true positives, *FP* the number of false positives, *TN* the number of true negatives, and *FN* the number of false negatives.

Sensitivity:

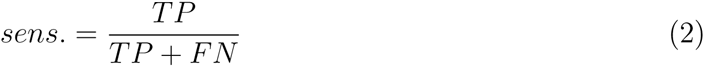

Specificity:

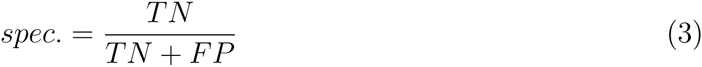

Balanced accuracy:

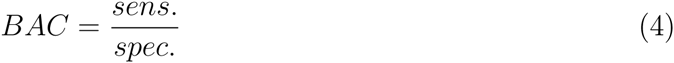

Matthews’ correlation coefficient:

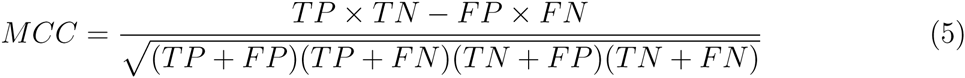

## Results and discussion

The sequence model was evaluated by its ability to correctly classify, on a per-kinase basis, kinase-specific phosphorylation sites out of the set of known kinase binding sites. Table 1 shows the averaged prediction accuracy for each of the kinase families; the full set of values are available in Table S-7 for human kinases, Table S-8 for mouse kinases, and Table S-9 for yeast kinases. The sequence model has good prediction accuracy over the kinases tested, with an average AUC of 0.79 across all human kinases. We found that 66% of kinases obtained an AUC of greater than 0.75, demonstrating that the model works well for the majority of kinases. We noticed particularly high accuracy for the CMGC kinases, where 17/20 of the kinases in this family obtained an AUC of greater than 0.8 (Table S-7); and also the atypical kinases, where all of those kinases obtained an AUC greater than 0.8, and 3/4 greater than 0.85 (Table S-7). The worst performing family appeared to be the tyrosine kinase family, where we found an average AUC of 0.62 – substantially lower than the overall average (of 0.79), and much lower than the accuracy from the various serine/threonine kinase families.

**Table 1:** Performance comparisons between predicting kinase-specific phosphorylation sites with a baseline model that only considers position-specific amino acid frequencies, and the sequence model. Results were generated using ten-fold cross-validation repeated across ten randomised data-set splits. Shown are the average and standard deviation of the AUC and AUC50 values.

| Family | AUC |  | AUC50 |  |
| --- | --- | --- | --- | --- |
|  | Baseline | Sequence | Baseline | Sequence |
| *Human* |  |  |  |  |
| CMGC | 0.80±0.013 | 0.84±0.014 | 0.08±0.013 | 0.21±0.026 |
| AGC | 0.76±0.017 | 0.79±0.018 | 0.15±0.028 | 0.21±0.029 |
| TK | 0.56±0.022 | 0.62±0.025 | 0.11±0.021 | 0.18±0.024 |
| CAMK | 0.73±0.023 | 0.77±0.024 | 0.11±0.014 | 0.19±0.027 |
| Other | 0.69±0.019 | 0.80±0.021 | 0.07±0.013 | 0.32±0.038 |
| STE | 0.71±0.031 | 0.79±0.052 | 0.23±0.049 | 0.38±0.053 |
| CK1 | 0.75±0.020 | 0.86±0.025 | 0.12±0.019 | 0.30±0.031 |
| Atypical | 0.84±0.009 | 0.87±0.008 | 0.18±0.008 | 0.20±0.030 |
| *Mouse* |  |  |  |  |
| CMGC | 0.74±0.016 | 0.79±0.016 | 0.14±0.017 | 0.24±0.029 |
| AGC | 0.72±0.025 | 0.75±0.032 | 0.17±0.034 | 0.26±0.051 |
| TK | 0.60±0.025 | 0.63±0.029 | 0.26±0.032 | 0.31±0.026 |
| *Yeast* |  |  |  |  |
| CMGC | 0.67±0.028 | 0.76±0.028 | 0.11±0.007 | 0.32±0.030 |
| AGC | 0.79±0.020 | 0.85±0.025 | 0.24±0.027 | 0.46±0.034 |
| CAMK | 0.64±0.024 | 0.78±0.024 | 0.05±0.017 | 0.34±0.037 |
| Other | 0.74±0.017 | 0.84±0.023 | 0.10±0.010 | 0.35±0.035 |

We compared the sequence model against a baseline model that only considered the position-specific amino acid frequencies. While the sequence model outperforms the baseline in general, we noticed that there was substantially higher accuracy at low false-positive levels as measured by the AUC50. In the “other” family of kinases, there was a greater than 3-fold increase in the AUC50, and in the CMGC and CK1 families we found a greater than 2-fold increase in AUC50.

On the mouse kinases, the model achieved a more moderate average AUC of 0.71, reflecting the diminished availability of positive training data when compared to human or yeast kinases. Similar to the results seen in the human kinases, however, the CMGC kinases performed the best, with an average AUC of 0.79, and the tyrosine kinases were again the worst performing, with an average AUC of 0.63.

The yeast kinase models performed quite well, achieving an average AUC of 0.81. In yeast, the best performing kinases were from the AGC family, with an average AUC of 0.85, and an AUC50 exceeding any other kinase family from mouse or human. We noticed that the sequence model had a substantial increase in accuracy when compared to the baseline – particularly at the low false-positive rates as measured by AUC50. The CAMK kinases recorded the sharpest increase, with an average AUC50 of over 6-fold greater than the baseline model. In general, we found that the use of k-mers offered a great advantage over the simpler representation of position-specific amino acid frequencies, and that this was particularly noticeable at low false-positive levels. Our results indicate that our combination of features offers a highly accurate model for predicting kinase phosphorylation sites across diverse kinase families and species.

In order to test whether sequence similarity within the phospho-peptides could be inflating prediction accuracy, we re-trained the sequence model on the similarity reduced data-set. Table S-10, Table S-11 and Table S-12 contain a comparison of the fully trained sequence model and the model trained on the reduced data-set. For the majority of kinases, the similarity reduction did not result in a decrease in AUC. On average, there was a negligible difference in AUC, with an average decrease across all kinases of 0.004 seen with the reduced data set. Similarly, differences in the average AUC50 were slight, and within the margin of error. This demonstrates that the prediction accuracy of the sequence model is not due to homologous phospho-peptides in the training data, and can be applied to unseen samples.

### Kinase substrate prediction

We compared the ability of the context model to predict kinase substrates against the combined (context plus sequence) model. The results summarised in Table 2 (see Table S-13, Table S-14 and Table S-15 for the complete set of kinases) demonstrate that across the kinase families, the incorporation of sequence data improved the ability of the model to predict kinase substrates. We noticed larger increases in prediction accuracy for the human CMGC, AGC and CAMK kinase families: the average AUC50 for CMGC increased from 0.31 to 0.43, AGC saw a similar increase from 0.21 to 0.34 and CAMK the largest – from 0.25 to 0.40.

**Table 2:** Performance comparisons between predicting kinase substrates with the context Bayesian network model, and with the combined sequence & context model. Results were generated using ten-fold cross-validation repeated across ten randomised data-set splits. Shown are the average and standard deviation of the AUC and AUC50 values.

|  | AUC |  | AUC50 |  |
| --- | --- | --- | --- | --- |
|  | Context | Combined | Context | Combined |
| *Human* |  |  |  |  |
| CMGC | 0.80±0.023 | 0.84±0.027 | 0.31±0.015 | 0.43±0.032 |
| AGC | 0.74±0.025 | 0.79±0.029 | 0.21±0.015 | 0.34±0.035 |
| TK | 0.81±0.027 | 0.82±0.026 | 0.31±0.020 | 0.39±0.039 |
| CAMK | 0.66±0.039 | 0.76±0.032 | 0.25±0.016 | 0.40±0.034 |
| Other | 0.80±0.034 | 0.81±0.037 | 0.36±0.029 | 0.47±0.044 |
| STE | 0.73±0.059 | 0.80±0.063 | 0.40±0.043 | 0.57±0.072 |
| CK1 | 0.79±0.035 | 0.81±0.028 | 0.39±0.032 | 0.41±0.042 |
| Atypical | 0.85±0.015 | 0.89±0.014 | 0.36±0.005 | 0.45±0.015 |
| *Mouse* |  |  |  |  |
| CMGC | 0.73±0.011 | 0.79±0.020 | 0.38±0.009 | 0.45±0.035 |
| AGC | 0.48±0.033 | 0.63±0.043 | 0.20±0.015 | 0.31±0.056 |
| TK | 0.61±0.045 | 0.78±0.052 | 0.25±0.020 | 0.46±0.052 |
| *Yeast* |  |  |  |  |
| CMGC | 0.65±0.032 | 0.76±0.042 | 0.22±0.020 | 0.44±0.050 |
| AGC | 0.57±0.043 | 0.71±0.048 | 0.26±0.036 | 0.48±0.048 |
| CAMK | 0.64±0.036 | 0.70±0.020 | 0.15±0.029 | 0.33±0.037 |
| Other | 0.60±0.036 | 0.75±0.045 | 0.21±0.019 | 0.40±0.033 |

While the context information accounts for the bulk of the accuracy, there were several examples of kinases where including the protein sequence in the model greatly improved prediction accuracy. In a few instances, prediction accuracy was increased from low or even random to a much higher value; for example the PKCI kinase improved from an AUC of 0.50 to an AUC of 0.77, and DYRK2 obtained a huge increase from an AUC of 0.63 to 0.91. There were also several examples of substantial accuracy gains, even when the kinase already had moderate to high accuracy in the context model; we observed that the prediction accuracy of GSK3A increased from 0.81 to 0.91, tyrosine kinase Syk increased from 0.81 to 0.90 and CAMK kinase Pim1 increased from 0.8 to 0.94. While there were examples of prediction accuracy decreasing when sequence information was added, these decreases were slight, indicating that the accuracy gains for incorporating sequence and context information far outweigh any potential losses.

In general, the accuracy for mouse kinases was more enhanced by the incorporation of sequence when compared to the accuracy for human kinases. We noticed that the accuracy for mouse AGC kinases was no greater than random for context alone, with a low AUC of 0.48. However, after the incorporation of sequence data, the AUC increased to a much higher value of 0.63. This is likely due to the size of the mouse protein-interactome, which is much smaller than the human version. The most substantial gains were made for the tyrosine kinases, where the average AUC for the family increase from 0.61 to 0.78 – a near 30% increase in prediction accuracy. There was a similar increase in the AUC50, from 0.25 to 0.46, indicating that the incorporation of the sequence model also made an important contribution at low false-positive levels.

The yeast kinases benefitted even more than the mouse kinases from the incorporation of sequence, with substantial increases to prediction accuracy observed across the four yeast kinase families. Prediction accuracy for yeast AGC and “other” kinases increased in AUC value by an average of 0.14 and 0.15 respectively, while CMGC kinases increased by an average of 0.09. We also found that the AUC50 increased by approximately two-fold for each of the four yeast kinase families. The results for mouse and yeast kinases indicate that the model is able to offset the reduced availability of the context information through the sequence data.

## Comparisons to alternative methods

We tested the ability of PhosphoPICK (i.e. the full PhosphoPICK workflow described in section “Prediction workflow”) to correctly classify the known kinase phosphorylation sites out of all potential sites within our set of phosphorylation substrates. Due to the number of potential phosphorylation sites (~170,000 S/T sites and ~30,000 Y sites), we tested prediction accuracy at more stringent levels of specificity – 99.9% and 99%. We compared the prediction sensitivity of PhosphoPICK with using sequence alone. We found that by combining the substrate score from the combined model with the site score from the sequence model, we were consistently able to improve prediction accuracy when compared to using the sequence model alone (Fig. 2).

**Figure 2:**
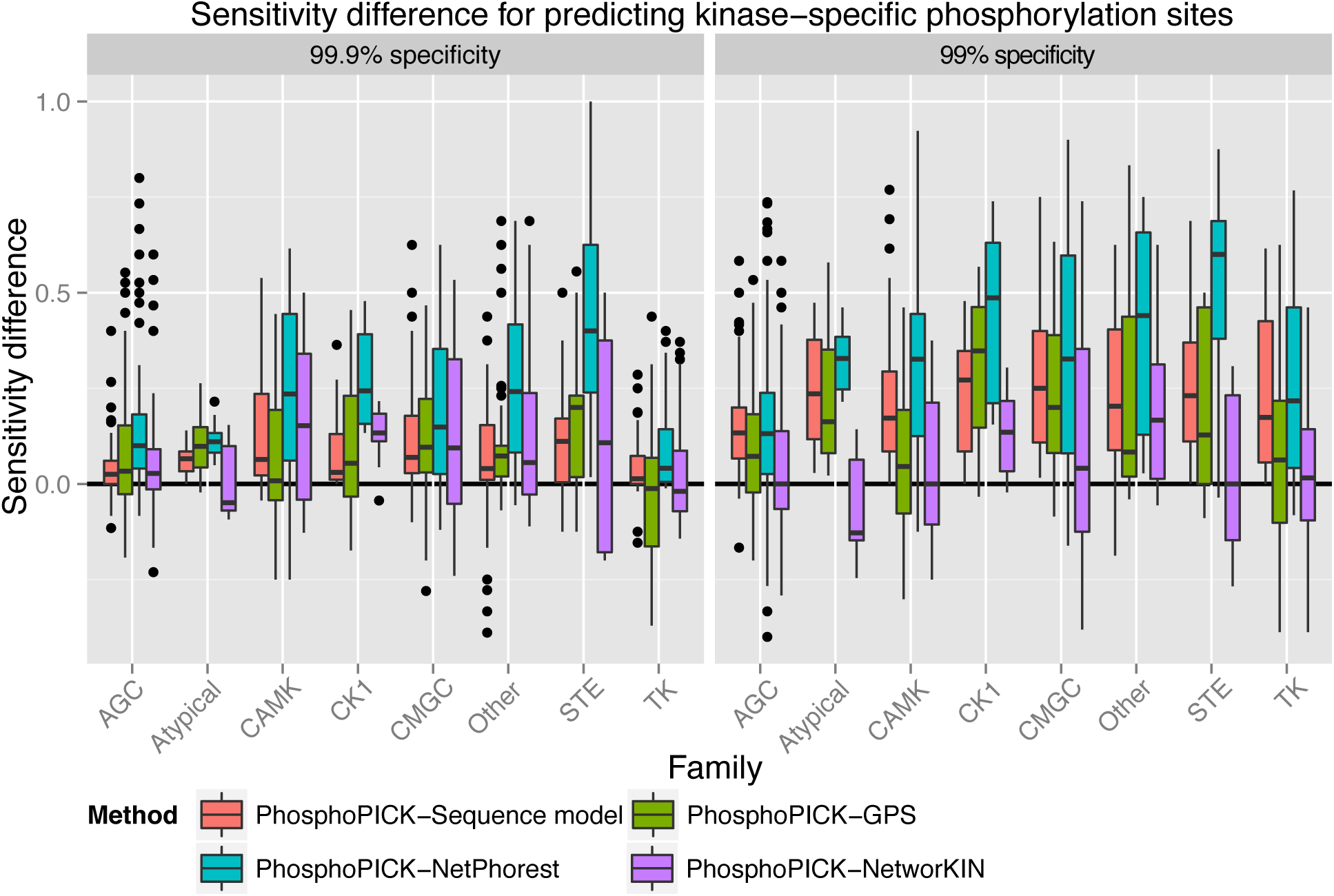
Sensitivity comparisons for predicting kinase-specific phosphorylation sites out of all potential phosphorylation sites in the protein training set between PhosphoPICK and alternative classification methods. Comparisons were made by performing cross-validation across ten data-set splits for each of the kinases. Sensitivity was calculated for all methods at two levels of specificity: 99.9% and 99%. Comparisons were made between PhosphoPICK and the sequence method alone, and between PhosphoPICK and three alternative predictors: GPS, NetPhorest and NetworKIN.

On average, the use of the combined model offered the greatest level of accuracy increase to kinases from the CMGC family, with an average sensitivity difference of 0.12 at 99.9% specificity and 0.27 at 99% specificity. This is consistent from our previous findings that the use of context offers greater support to phosphorylation site prediction from CMGC kinases. The CAMK kinases gained a similar level of sensitivity at the higher specificity threshold, though there was a smaller average sensitivity difference of 0.22 at the 99% specificity level. The AGC and TK kinases appeared to benefit the least, with a sensitivity difference at 99.9% specificity of 0.045 and 0.042, respectively.

We also compared the ability of PhosphoPICK to predict kinase-specific phosphorylation sites to three alternative methods: GPS 2.1,^13^ NetPhorest 2.0 and NetworKIN 3.0.^33^ We compared the prediction sensitivity of the different methods at the specificity levels described above. Fig. 2 shows the sensitivity difference between PhosphoPICK and the compared methods at two levels of specificity: 99.9% and 99%. The full set of comparisons for individual kinases are available in Table S-13 (comparisons at 99.9% specificity) and Table S-14 (comparisons at 99% specificity). We found that at the stricter level of specificity, PhosphoPICK obtained an increased level of sensitivity over the alternatives for most comparisons made. At the 99.9% specificity level, PhosphoPICK gained an average sensitivity increase of 9% when compared to NetworKIN, 10% compared to GPS and 22% compared to NetPhorest. At the 99% specificity level, PhosphoPICK gained average sensitivity increases of 6%, 18% and 35% when compared against NetworKIN, GPS and NetPhorest, respectively. There were some cases where PhosphoPICK performed worse than the alternatives – for example the tyrosine kinases, where we observed an average sensitivity difference against GPS of −0.014 at the 99.9% specificity level. We also noticed that PhosphoPICK performed worse on the atypical kinases when compared to NetworKIN, with a small difference in sensitivity at 99.9% specificity of −0.004, and a larger difference of −0.076 at 99% specificity.

## Evaluation using the hold-out set

PhosphoPICK contains the option to calculate P-values for predictions, representing the likelihood of obtaining a given prediction by chance, given how predictions are distributed over the proteome. To estimate the level of accuracy that is to be expected from using the fully trained model underlying the web-server, we evaluated prediction accuracy using our hold-out set of 145 substrates (of the kinases listed in Table 3) by calculating P-values of the predictions and considering predictions that fell below a P-value threshold of 0.005.

**Table 3:** Prediction accuracy on hold-out set for predicting kinase-specific phosphorylation sites (below a P-value threshold of 0.005) as measured by a variety of metrics – sensitivity, specificity, balanced accuracy (BAC) and Matthews’ correlation coefficient (MCC). Results were generated by training the model on the full training data set, and evaluating it on the hold-out set. Results represent the ability of PhosphoPICK to correctly predict the known kinase-specific phosphorylation sites out of all potential sites in the set of hold-out substrates. In total there were 14,617 S/T sites and 2,324 Y sites.

| Kinase | Positives | Sensitivity | Specificity | BAC | MCC |
| --- | --- | --- | --- | --- | --- |
| CDK2 | 72 | 0.36 | 0.96 | 0.66 | 0.12 |
| CDK1 | 39 | 0.51 | 0.93 | 0.72 | 0.09 |
| ERK2 | 55 | 0.22 | 0.98 | 0.60 | 0.08 |
| ERK1 | 56 | 0.29 | 0.98 | 0.63 | 0.12 |
| PKACA | 53 | 0.28 | 0.99 | 0.64 | 0.18 |
| PKCA | 40 | 0.15 | 0.97 | 0.56 | 0.04 |
| Akt1 | 15 | 0.4 | 0.98 | 0.69 | 0.09 |
| CK2A1 | 52 | 0.62 | 0.95 | 0.78 | 0.15 |
| Src | 34 | 0.03 | 0.99 | 0.51 | 0.02 |

We found that PhosphoPICK was generally able to maintain a high level of specificity, with an average specificity of 97% across the 9 kinases represented in the hold-out set (Table 3). There was a diverse range of sensitivity levels (from 3% for Src to 62% for CK2A1), with an average of 32% – well above what would be expected by chance given the percentage of false-positive predictions. This confident prediction accuracy on completely novel data indicates that PhosphoPICK is a reliable method for uncovering new kinase substrates and kinase-specific phosphorylation sites.

## Multiple kinases regulate nuclear localisation

We predicted NLSs using the NucImport predictor,^34^ a tool for predicting nuclear proteins and the location of their NLSs on the basis of protein interaction and sequence data (NucImport does not explicitly incorporate protein phosphorylation into its predictions). The complete human proteome (including isoforms) was run through NucImport and all proteins that were predicted to contain a type-1 classical NLS were retained – there were 4134 such proteins. The type-1 classical NLS contains an optimal four residue amino acid configuration of KR(K/R)R or K(K/R)RK.^35^ In order to investigate phosphorylation within a window surrounding the NLS, we defined a centre position, *P*_0_, as the third residue within the predicted NLS (in the literature, this position is usually designated “P4” ^22^), and cross-referenced the location of the signals with known phosphorylation sites from PhosphoSitePlus^®^. We identified 1,830 phosphorylation sites that were within a 20 residue window around *P*_0_. These phosphorylation sites were submitted to PhosphoPICK for analysis (predicting all human kinases), and a P-value threshold of 0.005 was used to return results with a high level of stringency.

In order to test for kinases that were regulating specific positions in relation to the NLS, we counted the number of predicted binding events for kinases at each position within the 20 residue window surrounding *P*_0_. To determine whether the number of predicted kinase binding sites near an NLS was greater than would be expected by chance, we tested for over-representation against all known phosphorylation sites within the set of predicted nuclear proteins. Over-representation was tested for using Fisher’s exact test with Bonferroni correction to obtain E-values (the P-values for the Fisher’s exact test were corrected by the total number of tests performed; i.e. the number of kinases multiplied by the number of sites – 2,247).

Fig. 3 shows the distribution of predicted binding sites for several kinases around the *P*_0_ position of the NLS. We found that there was higher phosphorylation activity downstream from the NLS, where protein kinase A (PKA), aurora kinase B (AurB), and Akt1 in particular were found to have the most significantly over-represented binding locations. At position 3 (*P*_3_), the most significant kinase was PKA (E = 2.03e^−38^), which was predicted to be phosphorylating 55/144 of the phosphorylation sites at that position. AurB had a pair of highly significant binding sites at positions 2 (E = 7.32e^−30^) and 3 (E = 2.4e^−21^).

**Figure 3:**
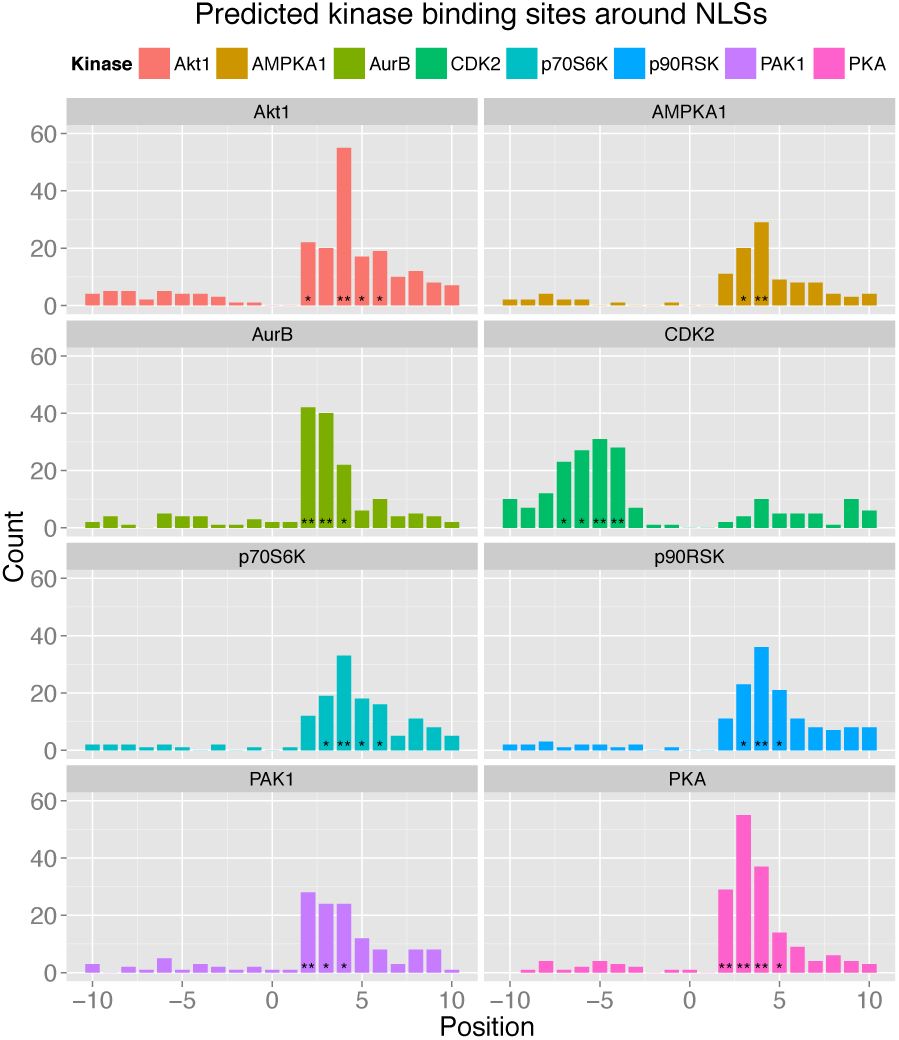
Distribution of predicted kinase phosphorylation sites surrounding NLSs.The locations of predicted NLSs were cross-referenced with phosphorylation sites from PhosphoSitePlus^®^ and PhosphoPICK was used to assign kinases to the sites. Count represents the number of times a kinase was predicted to phosphorylate a specific site relative to the NLS. Over-representation of a kinase for a particular site was assessed using a Fisher’s exact test with a Bonferroni multiple correction. (*) indicates an E-value < 0.05 and (**) an E-value < 1.0*E*^−10^.

There were fewer observations of kinases over-represented at phosphorylation sites upstream from the NLS, though we found that cyclic dependent kinase 2 (CDK2) and protein kinase C alpha (PKCa) were significantly over-represented at several upstream positions. At positions −4, −5 −6 and −7, CDK2 was found to have the most significant over-representation of sites compared to any other kinase. CDK2 was predicted to target 28/50 (E = 9.42e^−13^) of the phosphorylation sites at position −4, 31/61 (E = 2.1e^−13^) at position −5, 27/89 (E = 6.4e^−10^) at position −6 and 23/88 (E = 6.0e^−07^) at position −7.

To investigate whether the proteins being phosphorylated at these specific sites were involved in similar biological processes, we performed gene ontology (GO) term enrichment analyses. We performed the tests by taking a foreground set of proteins and testing for over-representation (Fisher’s exact test, with Bonferroni multiple correction) of terms in the foreground set against a background comprised of our set of phosphorylated nuclear proteins. Significant terms should therefore not simply represent general phosphorylation or nuclear functions, but functions specifically related to the kinase being tested.

We performed GO term enrichment tests on a kinase-specific basis, identifying substrates that were predicted to be phosphorylated within the 20 residue window surrounding *P*_0_. We also tested substrates that were predicted to be phosphorylated at the specific sites that were identified as being over-represented for the kinase being tested. We found that AurB substrates were enriched in the GO terms “chromosome”, “nucleosome” and “nucleosome assembly” (Table S-15). Interestingly, while the proteins phosphorylated by AurB at the *P*_3_ position were enriched in similar GO terms, the proteins phosphorylated at *P*_2_ returned no significant GO terms. While CDK2 substrates obtained the significant terms “chromosome”, “cell cycle”, “nucleus” and “DNA repair”, none of its significant binding site positions were found to be be associated with enriched GO terms (Table S-19).

We noticed that kinases with an over-representation of binding events at *P*_4_ consistently obtained a number of significant GO terms for substrates phosphorylated at that site. In addition to AurB mentioned above, PKA *P*_4_ substrates had 10 enriched GO terms (Table S-20), Akt1 had 4 (Table S-21), AMPKA1 and p70S6K both had 11 (Table-S22 and Table S-23, respectively) and p90RSK had 8 (Table S-24). We noticed that there was also some repetition of enriched GO terms among these kinases at *P* _4_ – the term for “fibroblast growth factor receptor (FGFR) signalling pathway” was the most significant *P*_4_ term for each of the AGC kinases (PKA, Akt1, p70S6K and p90RSK), and was the second most significant for AMPKA1 kinase. To determined whether phosphorylation at *P*_4_ in general was associated with specific functions (such as the FGFR signalling pathway) we did a GO term enrichment test with all substrates that were phosphorylated at that position, however no GO terms were found to be significant (Table S-25). This would indicate that the phosphorylation of the site at *P*_4_ does not by itself correspond to a particular function, rather this is dependent on the kinase regulating the site.

## Conclusions

The regulation of protein function through kinase-mediated phosphorylation is a complex process involving numerous aspects of cellular behaviour on the systems biology level, and the binding capacity of kinases to substrates on the molecular level. We have presented here a novel method for probabilistically modelling the sequence features that determine kinase binding at a molecular level. We have shown that PhosphoPICK is able to leverage these two diverse types of information and seamlessly integrate them into a model that can identify kinase substrates with high accuracy.

A benefit of the integration of sequence and context data into a single probabilistic model is the ability to take into account interdependance between these heterogeneous sources of information; i.e. the likelihood of seeing certain amino acids or k-mers in a protein may change depending on the context information, and similarly, the expectation of certain protein interactions can be influenced by the protein sequence. Indeed, we have found that the combined model can be used to query expected kinase binding sequence motifs and generate corresponding sequence logos^36^ based on context information presented to the model (see Methods S-1 for an example).

A counter-intuitive result seen as a part of the integration of sequence and context was that the performance seen in the sequence was not necessarily reflected in the combined model. The tyrosine kinases were a particularly interesting example; we found that while the tyrosine sequence models (for both human and mouse) were the least accurate amongst the sequence models, the mouse combined model benefited greatly from the incorporation of sequence, with a near two-fold increase seen in the AUC50. This is an indication that while the two individual systems – sequence and context – of predicting kinase binding events may be limited by themselves, the integration of the two can result in a much more powerful predictive model.

It was interesting to note that though the sequence model obtained the greatest accuracy (for phosphorylation site prediction) on the human kinases, the yeast kinases in general saw the highest increases in prediction accuracy (particularly as measured by AUC50) when the sequence model was incorporated into the context model. While the availability of context data (e.g. cell cycle data) is likely a factor in the observed differences in prediction performance between organisms, a uni-cellular organism like yeast would be expected to require less sophistication in the regulation of kinase activity than higher organisms. Consequently, the use of context factors is no doubt more important for understanding kinase targets in higher organisms.

For more complex organisms such as human and mouse, an additional realm of biology to consider in relation to phosphorylation and kinase activity is tissue and cell-type specificity. Protein phosphorylation has the potential to change substantially depending on the cell type, and the biological processes that kinases regulate can also vary depending on cell or tissue type. While there is limited amounts of consolidated tissue-specific phosphorylation data, there is growing amounts of tissue-specific protein expression data.^37^ In addition to protein expression data, the FANTOM consortium has profiled vast cell-type specific gene expression atlases. ^38^ Such data resources could make it possible to infer more probable candidate kinases based on which ones are available in the tissue or cell type of interest. While outside the scope of the current study, this would certainly make for an interesting avenue of exploration in future work.

A system-wide analysis of biological mechanisms has the potential to reveal functional trends that may not otherwise be apparent. Our analysis of the overlap of NLSs and phosphorylation events has shown that there are several kinases that may be implicated in the regulation of nuclear localisation through the phosphorylation of specific sites close to the NLS. Phosphorylation is a well-documented mechanism of nuclear localisation.^4,23,24,39—42^ Because classical NLSs are positively charged, introduction of a negatively charged phosphate group in the vicinity of the NLS would in general be expected to inhibit nuclear import, as previously demonstrated for CDK1-mediated phosphorylation at positions “P0” and “*P*-1” ^24^ (interestingly, these sites correspond to our *P*_–4_ and *P*_–5_ positions, which saw the most significant over-representation of CDK2 binding sites.). However, the effect will depend on the specific position that is phosphorylated, and in some positions phosphorylation can stimulate nuclear import.^4,23,39,40,42,43^

Several of the kinases identified in our study have previously been implicated in nuclear import. For example, the import of sex-determining factor SOX9 is regulated by PKA, whereby the phosphorylation of two phosphorylation sites (one next to the NLS) enhances SOX9 binding to importin *β*.^44^ Adenomatus polyposis coli (APC) is another example of a protein where nuclear import is regulated by phosphorylation. ^45^ In this case, APC contains two identified NLSs and a putative PKA-mediated phosphorylation site is positioned immediately after the second NLS, which leads to a reduction in APC nuclear localisation when the site is active. As a key regulator during mitosis, AurB is involved in several processes such as mitotic chromosome condensation, ^46^ and it has also been shown to phosphorylate residues within the vicinity of NLSs.^47^ The Akt kinase has been shown to be a regulator of nuclear localisation,^48^ and phosphorylation by Akt is able to impair the nuclear import of p27 *in vitro*.^49^ Similarly, CDK2 is known to be a regulator of nuclear localisation.^50^ While these studies confirm that these kinases are involved in nuclear localisation, our results shed light on specific mechanisms whereby nuclear localisation is controlled by the phosphorylation of key residues close to the NLS.

## Availability

PhosphoPICK is freely available online as a web-server, and can be used in two ways. A user can upload protein sequences, and select any number of kinases to obtain predictions for potential phosphorylation sites on the proteins. Significance of predictions can be gauged through the calculation of empirical P-values, and only results below a chosen level of significance returned. Visualisation of results is also available through a “Protein Viewer” page based on the BioJS^51^ package pViz.^52^ Secondly, the web-server allows for the construction of downloadable proteome-wide sets of kinase-substrate predictions for any of the kinases and species described in this paper. A more detailed description of the web-server workflow is available in Methods S-2.

## Acknowledgement

KALC was supported by the National Health and Medical Research Council (NHMRC) Career Development fellowship (APP1087415). BK is a NHMRC Senior Research Fellow (1003326 and 1110971).

## Supporting Information Available

**Table S-1**

Sequence model accuracy across human kinases when different percentages of kinase-substrate phosphorylation peptides were used to determine the set of k-mers added to the model. Table shows median AUC and AUC50 values for classifying kinase phosphorylation sites with the sequence model as determined by 10-fold cross-validation across 10 randomised data-set splits. Kinases are grouped according to their family, with the average prediction accuracy for each family shown.

**Table S-2**

Sequence model accuracy across mouse kinases when different percentages of kinase-substrate phosphorylation peptides were used to determine the set of k-mers added to the model. Table shows median AUC and AUC50 values for classifying kinase phosphorylation sites with the sequence model as determined by 10-fold cross-validation across 10 randomised data-set splits. Kinases are grouped according to their family, with the average prediction accuracy for each family shown.

**Table S-3**

Sequence model accuracy across yeast kinases when different percentages of kinase-substrate phosphorylation peptides were used to determine the set of k-mers added to the model. Table shows median AUC and AUC50 values for classifying kinase phosphorylation sites with the sequence model as determined by 10-fold cross-validation across 10 randomised data-set splits. Kinases are grouped according to their family, with the average prediction accuracy for each family shown.

**Table S-4**

Sequence model accuracy for varying window sizes in human kinases. Table shows accuracy values for classifying kinase phosphorylation sites with the sequence model as determined by 10-fold cross-validation across 10 randomised data-set splits. Prediction accuracy is shown using median and standard deviation of the AUC and AUC50 across the data-set splits. Varying window sizes were applied to determine the optimal window size on a kinase-specific basis. The window size determined for a kinase is highlighted through bold text. Optimal window size was determined primarily through AUC50 as a measure of the model’s accuracy at low false-positive rates. If accuracy did not increase through increasing window size, the lower window size was chosen. Kinases in the table are grouped according to family.

**Table S-5**

Sequence model accuracy for varying window sizes in mouse kinases. Table shows accuracy values for classifying kinase phosphorylation sites with the sequence model as determined by 10-fold cross-validation across 10 randomised data-set splits. Prediction accuracy is shown using median and standard deviation of the AUC and AUC50 across the data-set splits. Varying window sizes were applied to determine the optimal window size on a kinase-specific basis. The window size determined for a kinase is highlighted through bold text. Optimal window size was determined primarily through AUC50 as a measure of the model’s accuracy at low false-positive rates. If accuracy did not increase through increasing window size, the lower window size was chosen. Kinases in the table are grouped according to family.

**Table S-6**

Sequence model accuracy for varying window sizes in yeast kinases. Table shows accuracy values for classifying kinase phosphorylation sites with the sequence model as determined by 10-fold cross-validation across 10 randomised data-set splits. Prediction accuracy is shown using median and standard deviation of the AUC and AUC50 across the data-set splits. Varying window sizes were applied to determine the optimal window size on a kinase-specific basis. The window size determined for a kinase is highlighted through bold text. Optimal window size was determined primarily through AUC50 as a measure of the model’s accuracy at low false-positive rates. If accuracy did not increase through increasing window size, the lower window size was chosen. Kinases in the table are grouped according to family.

**Table S-7**

Comparison of prediction accuracy across human kinases between sequence model and baseline. Comparison of prediction accuracy across human kinases between predicting kinase-specific phosphorylation sites with a baseline model that only considers position-specific amino acid frequencies, and the sequence model. Kinases are grouped according to their family, with the average prediction accuracy for each family included. Results were generated using ten-fold cross-validation repeated across ten randomised data-set splits. Shown are the average and standard deviation of the AUC and AUC50 values.

**Table S-8**

Comparison of prediction accuracy across mouse kinases between sequence model and baseline. Comparison of prediction accuracy across mouse kinases between predicting kinase-specific phosphorylation sites with a baseline model that only considers position-specific amino acid frequencies, and the sequence model. Kinases are grouped according to their family, with the average prediction accuracy for each family included. Results were generated using ten-fold cross-validation repeated across ten randomised data-set splits. Shown are the average and standard deviation of the AUC and AUC50 values.

**Table S-9**

Comparison of prediction accuracy across yeast kinases between sequence model and baseline. Comparison of prediction accuracy across yeast kinases between predicting kinase-specific phosphorylation sites with a baseline model that only considers position-specific amino acid frequencies, and the sequence model. Kinases are grouped according to their family, with the average prediction accuracy for each family included. Results were generated using ten-fold cross-validation repeated across ten randomised data-set splits. Shown are the average and standard deviation of the AUC and AUC50 values.

**Table S-10**

Comparison of sequence model prediction accuracy across human kinases for training on full versus similarity-reduced data-sets. Comparison of prediction accuracy across human kinases between predicting kinase-specific phosphorylation sites using the sequence model trained on the full data-set, and when the model is trained on the similarity-reduced data-set. Prediction accuracy is calculated on the similarity-reduced data-set. If a kinase could not be trained on the reduced data-set due to too few positive training samples it was marked as “N/A”. Kinases are grouped according to their family, with the average prediction accuracy for each family included. Results were generated using ten-fold cross-validation repeated across ten randomised data-set splits. Shown are the average and standard deviation of the AUC and AUC50 values.

**Table S-11**

Comparison of sequence model prediction accuracy across mouse kinases for training on full versus similarity-reduced data-sets. Comparison of prediction accuracy across mouse kinases between predicting kinase-specific phosphorylation sites using the sequence model trained on the full data-set, and when the model is trained on the similarity-reduced data-set. Prediction accuracy is calculated on the similarity-reduced data-set. If a kinase could not be trained on the reduced data-set due to too few positive training samples it was marked as “N/A”. Kinases are grouped according to their family, with the average prediction accuracy for each family included. Results were generated using ten-fold cross-validation repeated across ten randomised data-set splits. Shown are the average and standard deviation of the AUC and AUC50 values.

**Table S-12**

Comparison of sequence model prediction accuracy across yeast kinases for training on full versus similarity-reduced data-sets. Comparison of prediction accuracy across mouse kinases between predicting kinase-specific phosphorylation sites using the sequence model trained on the full data-set, and when the model is trained on the similarity-reduced data-set. Prediction accuracy is calculated on the similarity-reduced data-set. If a kinase could not be trained on the reduced data-set due to too few positive training samples it was marked as “N/A”. Kinases are grouped according to their family, with the average prediction accuracy for each family included. Results were generated using ten-fold cross-validation repeated across ten randomised data-set splits. Shown are the average and standard deviation of the AUC and AUC50 values.

**Table S-13**

Combined model accuracy across human kinases compared to the context only model. Combined model accuracy across human kinases when compared to the context only model. Kinases are grouped according to their family, with the average prediction accuracy for each family included. Table shows accuracy values for classifying kinase substrates with both models as determined by 10-fold cross-validation across 10 randomised data-set splits. Prediction accuracy is shown using median and standard deviation of the AUC and AUC50 across the data-set splits.

**Table S-14**

Combined model accuracy across mouse kinases compared to the context only model. Combined model accuracy across mouse kinases when compared to the context only model. Kinases are grouped according to their family, with the average prediction accuracy for each family included. Table shows accuracy values for classifying kinase substrates with both models as determined by 10-fold cross-validation across 10 randomised data-set splits. Prediction accuracy is shown using median and standard deviation of the AUC and AUC50 across the data-set splits.

**Table S-15**

Combined model accuracy across yeast kinases compared to the context only model. Combined model accuracy across yeast kinases when compared to the context only model. Kinases are grouped according to their family, with the average prediction accuracy for each family included. Table shows accuracy values for classifying kinase substrates with both models as determined by 10-fold cross-validation across 10 randomised data-set splits. Prediction accuracy is shown using median and standard deviation of the AUC and AUC50 across the data-set splits.

**Table S-16**

Sensitivity differences for kinases at 99.9% specificity. Sensitivity differences for kinases at 99.9% specificity, where kinases are grouped according to their family, with the average sensitivity difference for each family included. The sensitivity difference between PhosphoPICK and each alternative method was measured for predicting kinase-specific phosphorylation sites out of all potential phosphorylation sites in our set of substrates. If we were unable to identify predictions for a kinase, it was marked as “N/A”.

**Table S-17**

Sensitivity differences for kinases at 99% specificity. Sensitivity differences for kinases at 99% specificity, where kinases are grouped according to their family, with the average sensitivity difference for each family included. The sensitivity difference between PhosphoPICK and each alternative method was measured for predicting kinase-specific phosphorylation sites out of all potential phosphorylation sites in our set of substrates. If we were unable to identify predictions for a kinase, it was marked as “N/A”.

**Table S-18**

Gene ontology (GO) term enrichment analysis for predicted AurB substrates. Shown are all positions that the kinase was found to be significantly over-represented at.

**Table S-19**

Gene ontology (GO) term enrichment analysis for predicted CDK2 substrates. Shown are all positions that the kinase was found to be significantly over-represented at.

**Table S-20**

Gene ontology (GO) term enrichment analysis for predicted PKA substrates. Shown are all positions that the kinase was found to be significantly over-represented at.

**Table S-21**

Gene ontology (GO) term enrichment analysis for predicted Akt1 substrates. Shown are all positions that the kinase was found to be significantly over-represented at.

**Table-S22**

Gene ontology (GO) term enrichment analysis for predicted AMPKA1 substrates. Shown are all positions that the kinase was found to be significantly over-represented at.

**Table S-23**

Gene ontology (GO) term enrichment analysis for predicted p70S6K substrates. Shown are all positions that the kinase was found to be significantly over-represented at.

Gene ontology (GO) term enrichment analysis for predicted p90RSK substrates. Shown are all positions that the kinase was found to be significantly over-represented at.

**Table S-25**

Gene ontology (GO) term enrichment analysis for substrates predicted to contain an NLS and a phosphorylation site at the specific position relative to the NLS. Table shows GO terms identified at each position in the 20-residue window surrounding the NLS.

**Methods S-1**

Identifying expected sequence motifs from context.

**Methods S-2**

Web-server workflow.

**S1 Fig**

Comparison of sequence logos for PKA kinase.

**Data S1**

Training data and model specification files for training the models presented in the paper. This material is available free of charge via the Internet at http://pubs.acs.org/.

## References

(1) Mayr, B.; Montminy, M. Nat. Rev. Mol. Cell. Biol. 2001, 2, 599–609.

(2) Liu, C.; Srihari, S.; Cao, K.-A. L.; Chenevix-Trench, G.; Simpson, P. T.; Ragan, M. A.; Khanna, K. K. Nucleic Acids Res. 2014, 42, 6106–6127.

(3) Dephoure, N.; Zhou, C.; Villén, J.; Beausoleil, S. A.; Bakalarski, C. E.; Elledge, S. J.; Gygi, S. P. Proc. Natl. Acad. Sci. U. S. A. 2008, 105, 10762–10767.

(4) Nardozzi, J.; Lott, K.; Cingolani, G. Cell Commun. Signaling 2010, 8, 32.

(5) Hornbeck, P. V.; Kornhauser, J. M.; Tkachev, S.; Zhang, B.; Skrzypek, E.; Murray, B.; Latham, V.; Sullivan, M. Nucleic Acids Res. 2012, 40, D261–D270.

(6) Hjerrild, M.; Gammeltoft, S. FEBS Lett. 2006, 580, 4764 – 4770.

(7) Hjerrild, M.; Stensballe, A.; Rasmussen, T.; Kofoed, C.; Blom, N.; Sicheritz-Ponten, T.; Larsen, M.; Brunak, S.; Jensen, O.; Ganuneltoft, S. J. Proteome Res. 2004, 3, 426–433.

(8) Manning, B.; Tee, A.; Logsdon, M.; Blenis, J.; Cantley, L. Mol. Cell 2002, 10, 151–162.

(9) Brinkworth, R. I.; Breinl, R. A.; Kobe, B. Proc. Natl. Acad. Sci. U. S. A. 2003, 100, 74–79.

(10) Kobe, B.; Kampmann, T.; Forwood, J. K.; Listwan, P.; Brinkworth, R. I. Biochim. Biophys. Acta 2005, 1754, 200 – 209.

(11) Wong, Y.-H.; Lee, T.-Y.; Liang, H.-K.; Huang, C.-M.; Wang, T.-Y.; Yang, Y.-H.; Chu, C.-H.; Huang, H.-D.; Ko, M.-T.; Hwang, J.-K. Nucleic Acids Res. 2007, 35, W588–W594.

(12) Ellis, J. J.; Kobe, B. PLoS ONE 2011, 6, e21169.

(13) Xue, Y.; Liu, Z.; Cao, J.; Ma, Q.; Gao, X.; Wang, Q.; Jin, C.; Zhou, Y.; Wen, L.; Ren, J. Protein Eng., Des. Sel. 2011, 24, 255–260.

(14) Blom, N.; Sicheritz-Pontén, T.; Gupta, R.; Gammeltoft, S.; Brunak, S. Proteomics 2004, 4, 1633–1649.

(15) Ingrell, C. R.; Miller, M. L.; Jensen, O. N.; Blom, N. Bioinformatics 2007, 23, 895–897.

(16) Zhu, G.; Liu, Y.; Shaw, S. Cell Cycle 2005, 4, 52 – 56.

(17) Good, M. C.; Zalatan, J. G.; Lim, W. A. Science 2011, 332, 680–686.

(18) Bloom, J.; Cross, F. R. Nat. Rev. Mol. Cell Biol. 2007, 8, 149–160.

(19) Ebisuya, M.; Kondoh, K.; Nishida, E. J. Cell Sci. 2005, 118, 2997–3002.

(20) Lapenna, S.; Giordano, A. Nat. Rev. Drug Discovery 2009, 8, 547–566.

(21) Patrick, R.; Lê Cao, K.-A.; Kobe, B.; Bodén, M. Bioinformatics 2015, 31, 382–389.

(22) Marfori, M.; Mynott, A.; Ellis, J. J.; Mehdi, A. M.; Saunders, N. F.; Curmi, P. M.; Forwood, J. K.; Bodén, M.; Kobe, B. Biochim. Biophys. Acta, Mol. Cell Res. 2011, 1813, 1562 – 1577.

(23) Christie, M.; Chang, C.-W.; Róna, G.; Smith, K. M.; Stewart, A. G.; Takeda, A. A.; Fontes, M. R.; Stewart, M.; Vértessy, B. G.; Forwood, J. K.; Kobe, B. J. Mol. Biol. 2015, –.

(24) Róna, G.; Borsos, M.; Ellis, J. J.; Mehdi, A. M.; Christie, M.; Környei, Z.; Neubrandt, M.; Tóth, J.; Bozóky, Z.; Buday, L.; Madarász, E.; Bodén, M.; Kobe, B.; Vértessy, B. G. Cell Cycle 2014, 13, 3551–3564.

(25) Stark, C.; Su, T.-C.; Breitkreutz, A.; Lourenco, P.; Dahabieh, M.; Breitkreutz, B.-J.; Tyers, M.; Sadowski, I. Database 2010, 2010.

(26) Chatr-aryamontri, A. et al. Nucleic Acids Res. 2013, 41, D816–D823.

(27) Franceschini, A.; Szklarczyk, D.; Frankild, S.; Kuhn, M.; Simonovic, M.; Roth, A.; Lin, J.; Minguez, P.; Bork, P.; von Mering, C.; Jensen, L. J. Nucleic Acids Res. 2013, 41, D808–D815.

(28) Olsen, J. V.; Vermeulen, M.; Santamaria, A.; Kumar, C.; Miller, M. L.; Jensen, L. J.; Gnad, F.; Cox, J.; Jensen, T. S.; Nigg, E. A.; Brunak, S.; Mann, M. Sci. Signal. 2010, 3, ra3.

(29) Camacho, C.; Coulouris, G.; Avagyan, V.; Ma, N.; Papadopoulos, J.; Bealer, K.; Madden, T. L. BMC Bioinf. 2009, 10, 1–9.

(30) Manning, G.; Whyte, D. B.; Martinez, R.; Hunter, T.; Sudarsanam, S. Science 2002, 298, 1912–1934.

(31) Do, C. B.; Batzoglou, S. Nat. Biotechnol. 2008, 26, 897–899.

(32) Baldi, P.; Brunak, S.; Chauvin, Y.; Anderson, C. A. F.; Nielsen, H. Bioinformatics 2000, 16, 412–424.

(33) Horn, H.; Schoof, E. M.; Kim, J.; Robin, X.; Miller, M. L.; Diella, F.; Palma, A.; Cesareni, G.; Jensen, L. J.; Linding, R. Nat. Meth. 2014, 11, 603–604.

(34) Mehdi, A.; Sehgai, M.; Kobe, B.; Bailey, T.; Bodén, M. Bioinformatics 2011, 27, 1239–1246.

(35) Kosugi, S.; Hasebe, M.; Matsumura, N.; Takashima, H.; Miyamoto-Sato, E.; Tomita, M.; Yanagawa, H. J. Biol. Chem. 2009, 284, 478–485.

(36) Crooks, G. E.; Hon, G.; Chandonia, J.-M.; Brenner, S. E. Genome Res. 2004, 14, 1188–1190.

(37) Uhlén, M. et al. Science 2015, 347.

(38) Consortium, T. F.; the RIKEN PMI,; (DGT), C. Nature 2014, 507, 462–470.

(39) Jans, D. A. Biochem. J. 1995, 311, 705–716.

(40) Jans, D. A.; Hubner, S. Physiol. Rev. 1996, 76, 651–685.

(41) Róna, G.; Marfori, M.; Borsos, M.; Scheer, I.; Takács, E.; Tóth, J.; Babos, F.; Magyar, A.; Erdei, A.; Bozóky, Z.; Buday, L.; Kobe, B.; Vértessy, B. G. Acta Crystallogr., Sect. D 2013, 69, 2495–2505.

(42) Fontes, M. R. M.; Teh, T.; Toth, G.; John, A.; Pavo, I.; Jans, D. A.; Kobe, B. Biochemical Journal 2003, 375, 339–349.

(43) Kosugi, S.; Hasebe, M.; Entani, T.; Takayama, S.; Tomita, M.; Yanagawa, H. Chem. Biol. 2008, 15, 940 – 949.

(44) Malki, S.; Nef, S.; Notarnicola, C.; Thevenet, L.; Gasca, S.; Mèjean, C.; Berta, P.; Poulat, F.; Boizet-Bonhourne, B. EMBO J. 2005, 24, 1798–1809.

(45) Zhang, F.; White, R. L.; Neufeld, K. L. Proc. Natl. Acad. Sci. U. S. A. 2000, 97, 12577–12582.

(46) Goldenson, B.; Crispino, J. D. Oncogene 2015, 34, 537–545.

(47) Guise, A. J.; Greco, T. M.; Zhang, I. Y.; Yu, F.; Cristea, I. M. Mol. Cell. Proteomics 2012, 11, 1220–1229.

(48) Biggs, W. H.; Meisenhelder, J.; Hunter, T.; Cavenee, W. K.; Arden, K. C. Proc. Natl. Acad. Sci. U. S. A. 1999, 96, 7421–7426.

(49) Liang, J.; Zubovitz, J.; Petrocelli, T.; Kotchetkov, R.; Connor, M. K.; Han, K.; Lee, J.H.; Ciarallo, S.; Catzavelos, C.; Beniston, R.; Franssen, E.; Slingerland, J. M. Nat. Med. 2002, 8, 1153–1160.

(50) Petersen, B. O.; Lukas, J.; Sørensen, C. S.; Bartek, J.; Helin, K. EMBO J. 1999, 18, 396–410.

(51) Gómez, J. et al. Bioinformatics 2013, 29, 1103–1104.

(52) Mukhyala, K.; Masselot, A. Bioinformatics 2014, 30, 3408–3409.

